# Construction of Discrete Element Model of Different Moisture Hygroscopic Fertilizer Particles and Fertilizer Discharging Verification

**DOI:** 10.1101/2024.09.03.611082

**Authors:** Chen Xiong Fei, Sun Ze Yu, Shi Yi Ze, Liu Mu Hua, Yu Jia Jia, Liu Jun An

## Abstract

The hygroscopic fertilizer particles were characterized based on discrete element simulation, and its accuracy was verified by the simulation of fertilizer discharge. Firstly, the repose angle of the fertilizer particles served as the key metric for assessment. A mathematical model characterizing the relationship between fertilizer moisture content and the repose angle was established, R^2^=0.9935; and a mathematical representation of the repose angle for fertilizer moisture content was established, R^2^=0.9933. Then, the moisture absorbing fertilizer particles was developed utilizing the integrated Hertz Mindlin with JKR model within EDEM system to account for particle adhesion; A robust correlation model between the repose angle of hygroscopic fertilizers and the discrete element method was established, p <0.0001; Finally, a correlation model between moisture content and significant parameters within the discrete element method was established, drawing upon the models for moisture content repose angle and repose angle discrete element parameters. The moisture-absorbing fertilizer particles model at moisture contents of 2% and 6% were resolved, with a relative error of the repose angle not exceeding 1.42% and a single circle fertilizer discharge error below 8.32%. The results show that the moisture content discrete element significant parameter correlation model can reliably and precisely forecast the discrete element parameters of various moisture content hygroscopic fertilizers. The hygroscopic fertilizer particle model can represent surface interaction characteristics of hygroscopic fertilizer granules. which has guiding significance for the design of precision fertilizer discharge technology device.

## 1. Introduction

Effective fertilization of farmland stands as a pivotal measure in guaranteeing consistent grain production yields, ensuring a stable yield of grain production is paramount, with mechanized precision fertilization emerging as a crucial method to achieve cost efficiency and optimal results. Creating a discrete element model for granular fertilizer is a crucial approach for conducting design optimization of essential equipment for precision fertilization, this method can significantly shorten the research and development cycle, lower experimentation costs, and enhance device performance [1-3]. Utilizing the discrete element method to analyze the intricate interaction between granular fertilizers, devices, and structures has emerged as a pivotal approach for investigating the kinematics and dynamics of agricultural granular materials [4,5]. This method is extensively applied in optimizing the design of fertilizer discharge structures [6-9].

Yuan Quanchun and Wang Xianliang [10-12] adjusted the discrete element parameters for organic fertilizer, soil, and composite fertilizer particles using the repose angle method for granular materials. Han Shujie and Wang Liming [13, 14] fine-tuned the discrete element parameters for organic fertilizer, soil, and fertilizer particles collectively using the repose angle method for granular materials. Zhu Xinhua [15] Bai, Jinyin [16] et al. developed a moisture content-repose angle model through the cylinder lifting method and regression fitting. Additionally, they formulated a discrete element model for sheep manure particles with varying moisture content by integrating Plackett-Burman and hill-climbing tests. Zhao Shuhong et al. [17] employed a discrete element model to study powdered organic fertilizers with varying moisture levels, aiming to develop an organic fertilizer feeding apparatus tailored for Northeast China. Yan Yinfa [8] et al. in their study, utilized the discrete element method to simulate and analyze the dynamics of motion and collisions among four different types of granular fertilizers namely nitrogen, phosphorus, potassium and organic fertilizers, in a four-fluted roller feed distributor. GELDART et al. [18] investigated the measurement technique for determining the angles of packing for eight different types of powdered particles. Sun J et al. [19] explored the dynamics of granular fertilizers within sheave-wheeled fertilizer dischargers and evaluated the characteristics of such dischargers. Their study involved a combined approach of simulation analysis and bench testing to investigate these features. The aforementioned research underscores the efficacy of employing the discrete element method to develop precise simulation models, offering a potent theoretical framework and efficient research approach for designing and examining pivotal components like fertilizer dischargers.

Contemporary literature demonstrates that numerous scholars, both domestically and internationally, have utilized discrete element analysis methods to investigate the kinematics and dynamics of fertilizer particles. They have applied these methods to design and optimize fertilizer discharger structures, aiming to enhance their performance further. Nevertheless, in practical production, the moisture absorption of fertilizer particles is inevitable, particularly in the high-temperature and high-humidity conditions prevalent in the southern rice region of China. This moisture uptake results in heightened surface adhesion and reduced mobility of the particles. Consequently, the fertilizer particles become more prone to adhering to the inner walls of the fertilizer discharging device, hindering proper discharge [20-22]. There remains a gap in the literature regarding the investigation of the kinematics and dynamics of viscous agricultural materials, such as granular fertilizers with varying moisture content post-absorption, utilizing the discrete element method. This knowledge deficiency underscores the need to address this issue, The objective of this study is to develop a correlation model between the moisture content of fertilizer and the repose angle using the cylinder lifting method and regression fitting. Utilizing Box-Behnken, Plackett-Burman, and steepest climb tests, a correlation model between the repose angle and discrete element parameters was established. Combining these equations yielded a predictive model for discrete element parameters of hygroscopic granular fertilizers with varying moisture levels. Subsequently, a cam top plate self-cleaning fertilizer feeding device was employed to simulate fertilizer discharge and conduct bench tests, validating the accuracy and reliability of the model for cohesive granular fertilizers with diverse moisture content. This study offers valuable research methodologies and theoretical frameworks for analyzing the motion characteristics of cohesive granular materials and designing precision fertilizer-feeding apparatuses.

## 2. Materials and Methods

### 2.1 External physical characteristics of moisture-absorbing granular fertilizers

Stanley compound fertilizer (17-17-17), a typical compound fertilizer in the southern rice region, was used as the test material. It was determined that, fertilizer moisture-absorbing granules’ moisture content in the southern rice region varied from 2% to 6%, with the sphericity φ being 0.887-0.899%, the equivalent diameter *d*_*e*_ being 3.27-3.53 mm, and the actual density being 1.622-1.766 g/cm^3^ [23].

### 2.2 Preparation of moisture-absorbing granular fertilizers with different moisture contents

In order to acquire hygroscopic and viscous granular fertilizers typical of the high-temperature and high-humidity conditions prevalent in the southern rice region, an MGC-450R artificial climate chamber was employed. This chamber allowed for the simulation of the specific climatic conditions of the southern region, enabling the preparation of moisture-absorbing viscous granular fertilizers with varying moisture levels. A plastic box measuring 52cm × 34cm, filled with a single layer of granular fertilizer, was positioned flat inside the artificial climate chamber for moisture absorption treatment. The temperature and humidity settings were adjusted to 25.00 ? and 80.00%, respectively.

The moisture content of fertilizer particles was measured by using Karl Fischer’s moisture titration method, with a target level of moisture content set at ± 0.5%, Figure 1 illustrates the variation in moisture content of moisture-absorbing fertilizer particles over time as they undergo moisture absorption The analysis of the moisture content-moisture absorption time curve revealed that the moisture content curve closely approximates a first-degree curve within the 0 to 8-hour timeframe. Consequently, moisture-absorbing granular fertilizers with moisture contents of 2%, 4%, and 6% were prepared accordingly, based on the moisture absorption time.

**Fig. 1.**
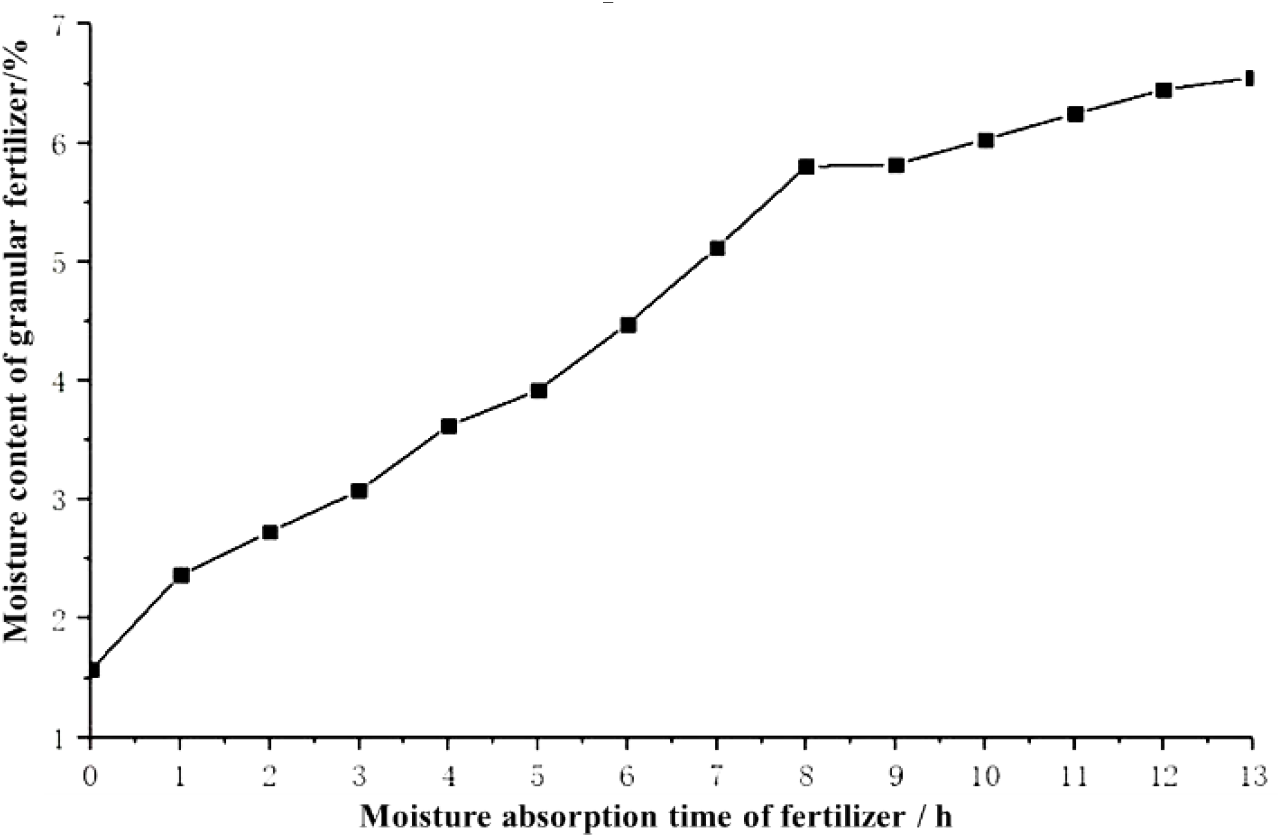
The plot illustrates the moisture content of fertilizer granules over time.

### 2.3 Modeling the correlation between moisture content and repose angle for hygroscopic and viscous granular fertilizers

The repose angle for granular fertilizers with varying moisture content was determined using the cylinder lifting method [24]. Figure 2 depicts the repose angle testing apparatus, comprising a bottomless cylinder (with an inner diameter of 32 mm and a height of 200 mm), a tension and compression testing machine (TMS-Pro texture instrument), a PC board platform, and a repose image capturing device’s angle.

**Fig. 2.**
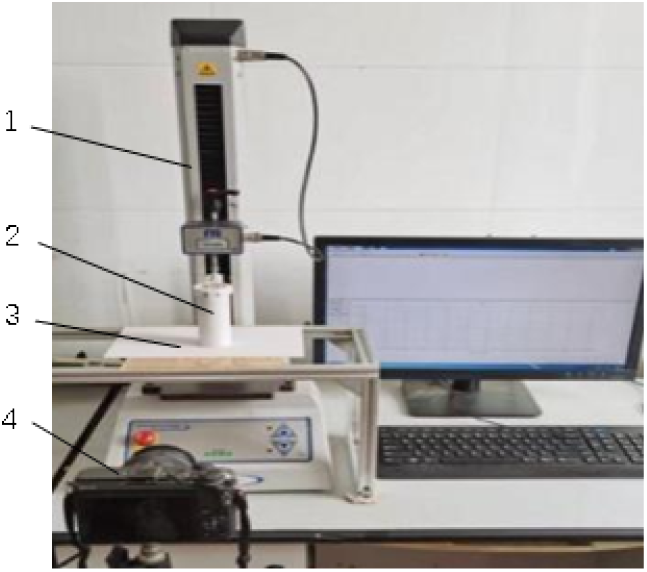
Apparatus used for angle of repose testing via the cylinder lifting method: 1. a tension and compression testing machine 2. PC bottomless cylinder 3. a PC board platform 4. an angle of rest image capturing device.

Prior to conducting the test, the cylinder was filled with moisture-absorbing granular fertilizers of varying moisture content. Subsequently, the tension and compression testing machine lifted the cylinder at a consistent speed, with the lifting rate set at 8.3 mm/s. The fertilizers spilled out from beneath the cylinder and accumulated on the PC board, forming a stable pile of fertilizer, as depicted in Figure 3. Images of the longitudinal section of the fertilizer pile were captured using a camera. The repose angle of the fertilizer was determined through the image-based digital simulation method [12, 16, 25]. The test was conducted ten times for fertilizers with different moisture contents to obtain the average value. A mathematical model correlating moisture content and repose angle was developed utilizing an Excel data analysis tool.

**Fig. 3.**
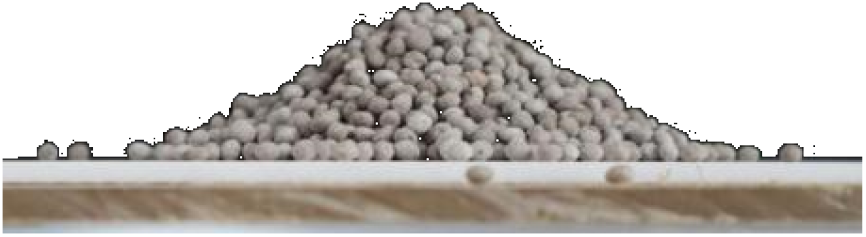
Schematic representation of granular fertilizer arrangement on the PC board.

**Fig. 4.**
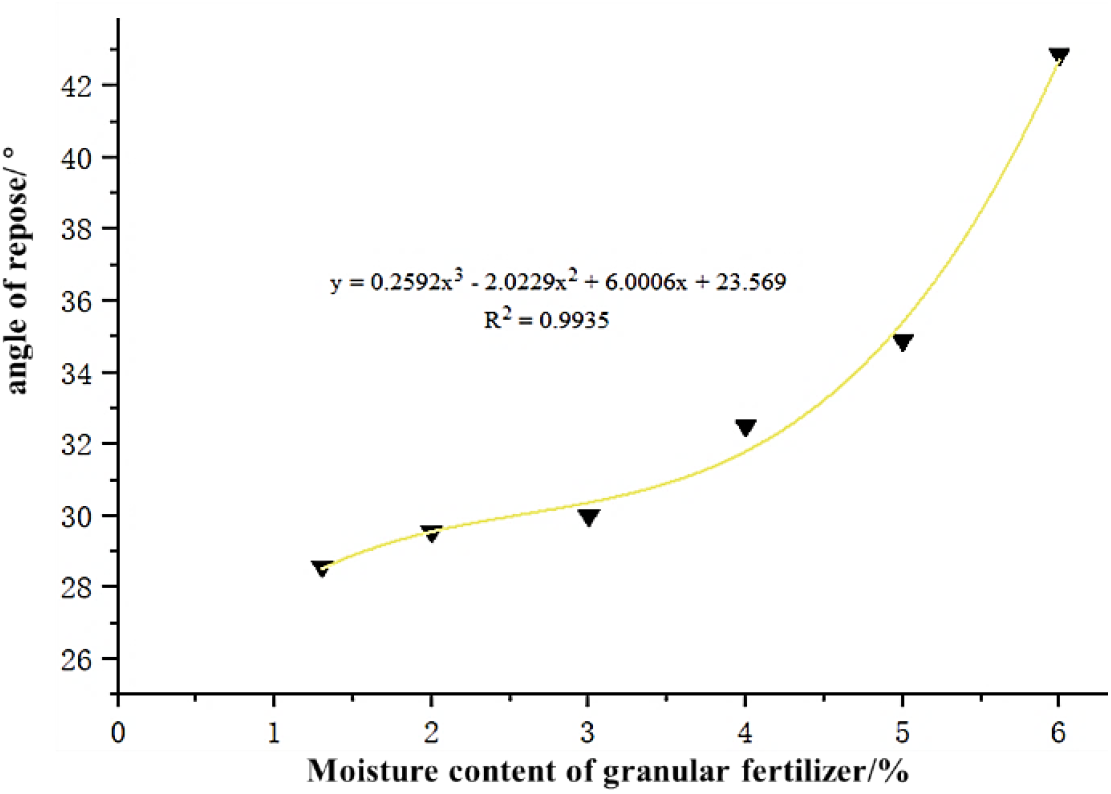
Fertilizer moisture content and angle of repose.

### 2.4 Discrete elemental modeling of hygroscopic and viscous granular fertilizers

#### 2.4.1 Intrinsic model parameters

Based on previous research, the observed variation in water content of moisture-absorbing viscous granular fertilizers in the southern rice region ranged from 2% to 6% [23]. Following the research method outlined in reference [13,14], moisture-absorbing viscous granular fertilizers with a moisture content of 4% were initially chosen to establish a discrete element model. These granules were assigned an equivalent diameter of 3.43 mm and a density of 1667 kg/m3. The intrinsic parameters of the fertilizer and PC board (fertilizer feeding material) are presented in Table 1.

**Table 1.** Intrinsic parameters of fertilizers and PC.

| Materials | Density g/cm<sup>3</sup> | Poisson ratio | Shear modulus/pa |
| --- | --- | --- | --- |
| Fertilizers | 1.662~1.766 | 0.2~0.5 | $1 \times 10^7 \sim 3.6 \times 10^7$ |
| PC board | 1.18 | 0.39 | $9 \times 10^9$ |

#### 2.4.2 Contact model parameters

Considering characteristics of moisture-absorbing viscous granular fertilizers, characterized by poor mobility and large adhesion force, contact model selected for moisture-absorbing granular fertilizers within the EDEM system is the “Hertz-Mindlin with JKR” model, abbreviated as the JKR model. The JKR model is particularly suitable for capturing adhesion and agglomeration of granular materials arising from factors such as electrostatic force between particles, moisture content and other reasons [26-28]. The contact parameters and the range of values of JKR model parameters were established by integrating data from Generic EDEM material model database (GEMM database) with intrinsic parameters of fertilizer and PC board, as outlined in Table 2.

**Table 2.** Fertilizer and PC contact parameters, JKR model parameter range.

| Parameters | Values |
| --- | --- |
| Static friction coefficient of fertilizer-fertilizer | 0.2~1.5 |
| Dynamic friction coefficient of fertilizer-fertilizer | 0.4~1 |
| Recover coefficient of fertilizer-fertilizer | 0~0.5 |
| JKR surface energy of fertilizer-fertilizer, J/m<sup>2</sup> | 0~2 |
| Static friction coefficient of fertilizer-pc | 0.2~0.6 |
| Dynamic friction coefficient of fertilizer-pc | 0.2~0.5 |
| Recover coefficient of fertilizer-pc | 0.1~0.5 |

### 2.5 Construction of the angle of repose-discrete element parameter model for moisture-absorbing viscous granular fertilizers

#### 2.5.1 Simulation of angle of rest of moisture-absorbing viscous granular fertilizers

To ensure consistency with the physical test, the simulation process was configured accordingly. A 3D model of the cylinder and PC board, created at a 1:1 scale using SolidWorks 3D modeling software, was imported into the EDEM simulation system in STEP format. The simulation was saved at intervals of 0.05 seconds, with a time step set to 20% and a grid size three times larger than the size of the smallest particle entity. A model consisting of a cylinder filled with 3000 particles of hygroscopic granular fertilizer was utilized. The cylinder ascended at a constant speed of 8.3 mm/s for a duration of 40 seconds until the kinetic energy of the particles reached zero. Following the simulation’s conclusion, the MATLAB image processing method was employed to extract the repose angle of the fertilizers in both the XOZ plane and the YOZ plane, respectively [12], test results were obtained by averaging the values from three repeated tests.

#### 2.5.2 Construction of a model correlating repose angle with discrete element parameters moisture-absorbing viscous granular fertilizers

The Plackett-Burman experimental design was conducted using Design Expert software, Using the repose angle of the fertilizer as the response variable, significant effects on the repose angle were screened from 10 actual parameters and 1 virtual parameter using Design Expert software. Each parameter was assigned two levels, high and low, which were coded as +1 and -1, respectively, as outlined in Table 3. Thirteen experimental simulations with varying treatments were conducted to determine the repose angle of the fertilizer. The experimental scheme and results are presented in Section 2.2.

**Table 3.** List of Placktt-Burman test parameters.

| Number | Parameters | Low-level (-1) | High-level(+1) |
| --- | --- | --- | --- |
| A | Poisson ratio of fertilizer | 0.2 | 0.5 |
| B | Shear modulus of fertilizer, Pa | 1x107 | 3.6x107 |
| C | Static friction coefficient of fertilizer-fertilizer | 0.2 | 1.5 |
| D | Dynamic friction coefficient of fertilizer-fertilizer | 0.4 | 1 |
| E | Recover coefficient of fertilizer-fertilizer | 0 | 0.5 |
| F | JKR surface energy of fertilizer-fertilizer, J/m2 | 0 | 2 |
| G | Static friction coefficient of fertilizer-pc | 0.2 | 0.6 |
| H | Dynamic friction coefficient of fertilizer-pc | 0.2 | 0.5 |
| I | Recover coefficient of fertilizer-pc | 0.1 | 0.5 |
| J | Virtual parameter | -1 | 1 |
| K | Virtual parameter | -1 | 1 |

To narrow the parameter range and determine the optimal interval, the three parameters with relatively strong influence were increased by equal gradient based on the Placktt-Burman test results. Furthermore, the average values obtained from the physical tests were utilized as the initial parameters for the hygroscopic granular fertilizer model. A steepest hill climbing test was then conducted to determine the range of values for the significant parameters. The protocol and results of the climb test are presented in section 2.3.

Following the results of the hill climbing test, a Box-Behnken test was designed to conduct the response surface simulation test for the repose angle. This allowed for the extraction of the repose angle for each treatment, The test scheme and results, including the level parameters, are presented in Section 2.4, as outlined in Table 4.

**Table 4.** Table of Box-Behnken significance level parameters.

| Level | Poisson ratio of fertilizer B | Surface energy of fertilizer F | Recover coefficient of fertilizer-pc I |
| --- | --- | --- | --- |
| -1 | $1 \times 10^7$ | 0 | 0.1 |
| 0 | $1.65 \times 10^7$ | 0.5 | 0.2 |
| 1 | $2.3 \times 10^7$ | 1 | 0.3 |

Using the repose angle corresponding to three types of moisture-absorbent granular fertilizers as the target variable, repose angle-discrete element parameter model obtained from the Box-Behnken test was solved to determine the optimal combination of discrete element parameters. Utilizing the optimal combination of discrete element parameters (with the values of non-significant parameters consistent with those determined in the hill climbing test), The simulation of cylinder lifting was conducted in the EDEM system to measure the repose angle. The error between the simulated repose angle and the actual value was calculated to verify the accuracy of the discrete element parameter model for moisture-absorbing granular fertilizers.

## 3. Results and Analysis

### 3.1 Modeling the correlation between moisture content and angle of repose

Table 5 presents the results of the repose angle for granular fertilizer with varying moisture content, as measured by the cylinder lifting test, a polynomial regression was fitted to the data of moisture content and repose angle of the granular fertilizer, resulting in the moisture content-repose angle model (as shown in Equation 1).

**Table 5.** Table of Box-Behnken significance level parameters.

| Moisture content of granular fertilizer/% | Angle of rest/° |
| --- | --- |
| 2 | 29.54 |
| 3 | 30.00 |
| 4 | 32.50 |
| 5 | 34.89 |
| 6 | 42.87 |

A polynomial regression analysis was conducted on the data of moisture content and repose angle of the granular fertilizer, yielding the moisture content-repose angle model, The correlation coefficient of the model was determined to be 0.9935. The repose angle of the granular fertilizer increases with the rise in moisture content, attributable to the decreased mobility and increased adhesion of the granular fertilizer following hygroscopicity.

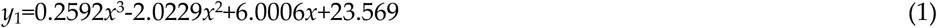

Where x represents moisture content, %; y represents the angle of rest, °.

### 3.2 Results and analysis screening significant parameters in discrete element modeling

Design Expert software was employed to conduct the Plackett-Burman experimental design. The experimental design and results are presented in Table 6, where A-J represent the coding values for the factors. The significance analysis of the test results, prioritizing parameters, reveals that the surface energy of fertilizer-fertilizer (F), the coefficient of restitution of fertilizer-PC (I), and the shear modulus of fertilizer (B) are particularly influential.

**Table 6.**
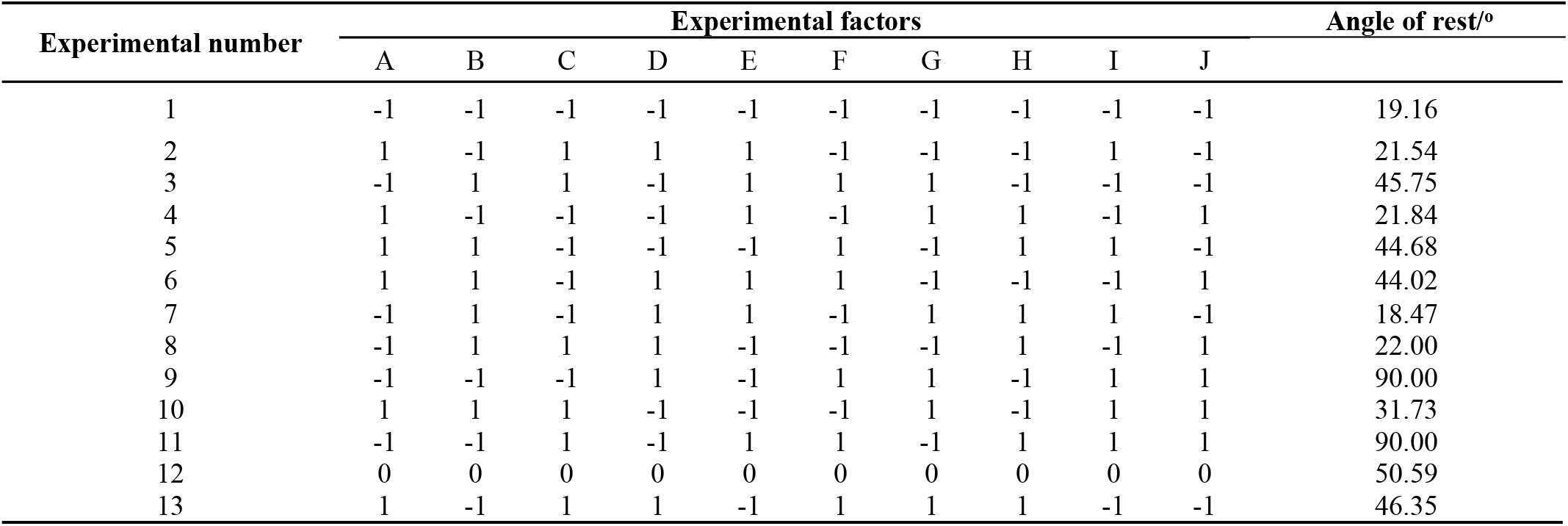
Placktt-Burman experimental design and results.

| Experimental number | Experimental factors |  |  |  |  |  |  |  |  |  | Angle of rest/° |
| --- | --- | --- | --- | --- | --- | --- | --- | --- | --- | --- | --- |
|  | A | B | C | D | E | F | G | H | I | J |  |
| 1 | -1 | -1 | -1 | -1 | -1 | -1 | -1 | -1 | -1 | -1 | 19.16 |
| 2 | 1 | -1 | 1 | 1 | 1 | -1 | -1 | -1 | 1 | -1 | 21.54 |
| 3 | -1 | 1 | 1 | -1 | 1 | 1 | 1 | -1 | -1 | -1 | 45.75 |
| 4 | 1 | -1 | -1 | -1 | 1 | -1 | 1 | 1 | -1 | 1 | 21.84 |
| 5 | 1 | 1 | -1 | -1 | -1 | 1 | -1 | 1 | 1 | -1 | 44.68 |
| 6 | 1 | 1 | -1 | 1 | 1 | 1 | -1 | -1 | -1 | 1 | 44.02 |
| 7 | -1 | 1 | -1 | 1 | 1 | -1 | 1 | 1 | 1 | -1 | 18.47 |
| 8 | -1 | 1 | 1 | 1 | -1 | -1 | -1 | 1 | -1 | 1 | 22.00 |
| 9 | -1 | -1 | -1 | 1 | -1 | 1 | 1 | -1 | 1 | 1 | 90.00 |
| 10 | 1 | 1 | 1 | -1 | -1 | -1 | 1 | -1 | 1 | 1 | 31.73 |
| 11 | -1 | -1 | 1 | -1 | 1 | 1 | -1 | 1 | 1 | 1 | 90.00 |
| 12 | 0 | 0 | 0 | 0 | 0 | 0 | 0 | 0 | 0 | 0 | 50.59 |
| 13 | 1 | -1 | 1 | 1 | -1 | 1 | 1 | 1 | -1 | -1 | 46.35 |

**Table 7.**
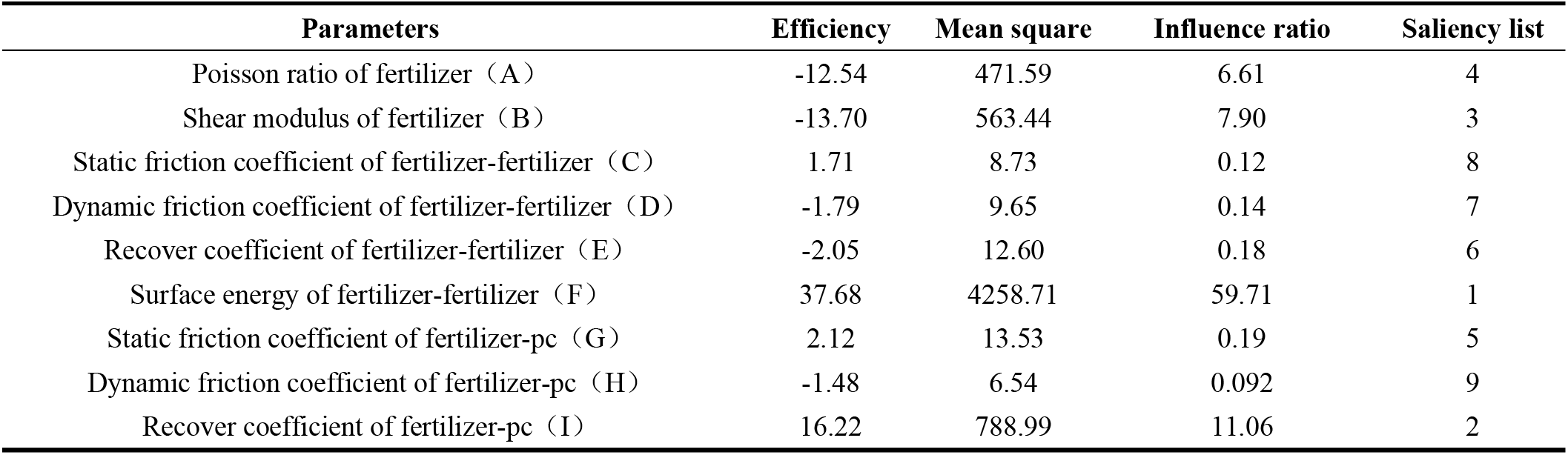
Placktt-Burman test parameter significance analysis.

| Parameters | Efficiency | Mean square | Influence ratio | Saliency list |
| --- | --- | --- | --- | --- |
| Poisson ratio of fertilizer (A) | -12.54 | 471.59 | 6.61 | 4 |
| Shear modulus of fertilizer (B) | -13.70 | 563.44 | 7.90 | 3 |
| Static friction coefficient of fertilizer-fertilizer (C) | 1.71 | 8.73 | 0.12 | 8 |
| Dynamic friction coefficient of fertilizer-fertilizer (D) | -1.79 | 9.65 | 0.14 | 7 |
| Recover coefficient of fertilizer-fertilizer (E) | -2.05 | 12.60 | 0.18 | 6 |
| Surface energy of fertilizer-fertilizer (F) | 37.68 | 4258.71 | 59.71 | 1 |
| Static friction coefficient of fertilizer-pc (G) | 2.12 | 13.53 | 0.19 | 5 |
| Dynamic friction coefficient of fertilizer-pc (H) | -1.48 | 6.54 | 0.092 | 9 |
| Recover coefficient of fertilizer-pc (I) | 16.22 | 788.99 | 11.06 | 2 |

### 3.3 Establishment of optimal intervals for significant parameters in discrete element modeling

The specific design of parameter levels and the results of the steepest hill climbing test are presented in Table 8. As the parameter increment increases, the relative error in the repose angle gradually rises from a negative value, and the parameter gradient increases from No. 1 to No. 2, The relative error transitions from a negative to a positive value, and the absolute value of the relative error for the No. 2 test is the smallest. Hence, it can be inferred that the parameter region adjacent to No. 2 is optimal. This explains why the parameter for the No. 2 test was selected as the center point, Consequently, No. 3 and No. 1 were designated as the positive and negative levels, respectively, to conduct the response surface simulation for repose angle. This facilitated the extraction of repose angle for each treatment group, in accordance with the Box-Behnken experimental design.

**Table 8.**
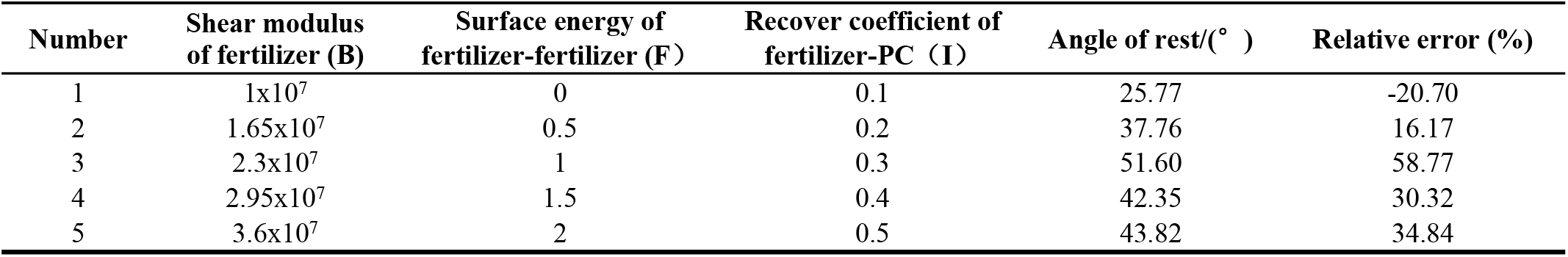
Steepest climb test design and results.

| Number | Shear modulus of fertilizer (B) | Surface energy of fertilizer-fertilizer (F) | Recover coefficient of fertilizer-PC (I) | Angle of rest/(°) | Relative error (%) |
| --- | --- | --- | --- | --- | --- |
| 1 | 1x10<sup>7</sup> | 0 | 0.1 | 25.77 | -20.70 |
| 2 | 1.65x10<sup>7</sup> | 0.5 | 0.2 | 37.76 | 16.17 |
| 3 | 2.3x10<sup>7</sup> | 1 | 0.3 | 51.60 | 58.77 |
| 4 | 2.95x10<sup>7</sup> | 1.5 | 0.4 | 42.35 | 30.32 |
| 5 | 3.6x10<sup>7</sup> | 2 | 0.5 | 43.82 | 34.84 |

### 3.4 Mathematical modeling of the correlation between angle of rest-discrete element parameter correlation of moisture-absorbing viscous granular fertilizers

The parameter identified as No. 2 in the steepest hill climbing test was designated as the 0 level, The parameter labeled as No. 1 was assigned the -1 level, and the No. 3 parameter was assigned the +1 level. The remaining parameter values remained consistent with those used in the hill climbing test [29]. The parameters for each level and the corresponding test results are displayed in Table 9.

**Table 9.** Box-Behnken experimental design and its results.

| Number | Shear modulus of fertilizer B | Surface energy of fertilizer F | Recover coefficient of fertilizer-PCI | Angle of rest/° |
| --- | --- | --- | --- | --- |
| 1 | 1.65x10<sup>7</sup> | 0.5 | 0.2 | 38.865 |
| 2 | 2.30x10<sup>7</sup> | 0 | 0.2 | 22.671 |
| 3 | 2.30x10<sup>7</sup> | 0.5 | 0.1 | 37.110 |
| 4 | 1.65x10<sup>7</sup> | 1 | 0.3 | 50.324 |
| 5 | 1.65x10<sup>7</sup> | 0.5 | 0.2 | 38.865 |
| 6 | 1.00x10<sup>7</sup> | 0 | 0.2 | 27.042 |
| 7 | 1.65x10<sup>7</sup> | 0.5 | 0.2 | 38.865 |
| 8 | 2.30x10<sup>7</sup> | 0.5 | 0.3 | 36.818 |
| 9 | 1.00x10<sup>7</sup> | 0.5 | 0.1 | 39.376 |
| 10 | 1.00x10<sup>7</sup> | 1 | 0.2 | 46.242 |
| 11 | 2.30x10<sup>7</sup> | 1 | 0.2 | 49.047 |
| 12 | 1.65x10<sup>7</sup> | 0.5 | 0.2 | 38.865 |
| 13 | 1.65x10<sup>7</sup> | 1 | 0.1 | 47.319 |
| 14 | 1.65x10<sup>7</sup> | 0.5 | 0.2 | 38.865 |
| 15 | 1.65x10<sup>7</sup> | 0 | 0.3 | 22.548 |
| 16 | 1.65x10<sup>7</sup> | 0 | 0.1 | 22.740 |
| 17 | 1.00x10<sup>7</sup> | 0.5 | 0.3 | 40.260 |

**Table 10.** Box-Behnken test quadratic polynomial regression model analysis of variance.

| Source of variance | Sum of squares | Degree of freedom | Mean square | F value | p value |
| --- | --- | --- | --- | --- | --- |
| Model | 1053.71 | 9 | 139.30 | 114.64 | < 0.0001** |
| B-Shear modulus of fertilizer | 6.61 | 1 | 6.61 | 5.44 | 0.0524 |
| F-Surface energy of fertilizer | 1198.81 | 1 | 1198.81 | 986.62 | < 0.0001** |
| I-Static friction coefficient of fertilizer-PC | 1.45 | 1 | 1.45 | 1.19 | 0.3109 |
| BF | 12.87 | 1 | 12.87 | 10.60 | 0.0140* |
| BI | 0.35 | 1 | 0.35 | 0.28 | 0.6102 |
| FI | 2.56 | 1 | 2.56 | 2.10 | 0.1903 |
| B<sup>2</sup> | 2.015E-003 | 1 | 2.015E-003 | 1.658E-003 | 0.9687 |
| F<sup>2</sup> | 29.27 | 1 | 29.27 | 24.09 | 0.0017** |
| I<sup>2</sup> | 1.04 | 1 | 1.04 | 0.85 | 0.3867 |
| Residual | 8.51 | 7 | 1.22 |  |  |
| Lack of Fit | 8.51 | 3 | 2.84 |  |  |
| Pure Error | 0.000 | 4 |  |  |  |
| Cor Total | 1262.22 | 16 |  |  |  |

$R^2 = 0.9933$ ; $R^2_{adj} = 0.9846$ ; CV=2.95%; Adeq Precisor=33.203

A diverse regression analysis of the Box-Behnken test parameters was conducted to derive the quadratic polynomial fitting equation for the correlation model between the repose angle (*y*_*2*_) and the discrete element parameters, given by:

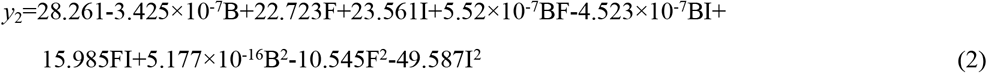

The variance analysis of the regression model was conducted, resulting in a p-value <0.0001. Decision coefficient R^2^=0.9933, while the adjusted coefficient R^2^_adj_ =0.9846. Both coefficients are close to 1, with a difference of less than 0.2, indicating well-fitted regression equation. Precision Adeq Precisior=33.203>4 indicates the reasonableness of the model. F and F^2^ exhibit a highly significant effect on the repose angle (p<0.01), while parameters B, I, BF, FI and I^2^ demonstrate a significant effect on the repose angle (0.1<p<0.5).

### 3.5 Prediction of angle of repose based on discrete element parameters for fertilizers with varying moisture content of hygroscopic particles

The repose angle of moisture-absorbing granular fertilizers with water content of 2%, 4%, and 6% was considered as the target value to solve the regression model. This facilitated the prediction of optimal combinations of parameter values for the shear modulus (B) of the three groups of fertilizers, fertilizer surface energy (F), and coefficient of restitution of fertilizer-PC-board (I). The optimal combination of fertilizer intrinsic contact parameters was utilized to conduct cylinder lifting simulations, resulting in the determination of the optimal parameter combination for the hygroscopic granular fertilizer model at each moisture content. The minimum value of relative error of the repose angle served as the boundary condition, with the results shown in Table 11.

**Table 11.**
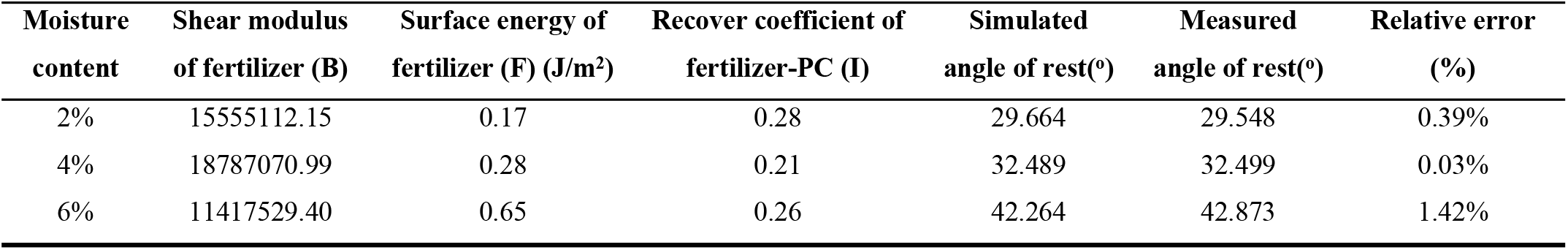
Rest angle results in simulations with different solution parameters for fertilizers with different moisture contents.

Figure 5 illustrates the simulated and measured repose angle for moisture-absorbing viscous granular fertilizers with 2%, 4%, and 6% water content. It indicates that the viscosity characteristics of moisture-absorbing viscous granular fertilizers increase with the rise in moisture content. Fertilizers with 2% and 4% moisture content exhibit less viscosity, while granules with 6% moisture content display significant viscous cohesion. As indicated in Table 11, the relative errors between the modeled repose angle and the actual repose angle for viscous granular fertilizers with 2% and 6% moisture content were 0.39% and 1.42%, respectively. Moreover, the shapes of the repose angle for the three models of moisture content viscous fertilizers were essentially identical to the actual shapes. The findings indicate that the model, which considers fertilizer moisture content as a significant parameter, accurately forecasts the discrete element parameters of hygroscopic granular fertilizers across different levels of moisture content.

**Fig. 5.**
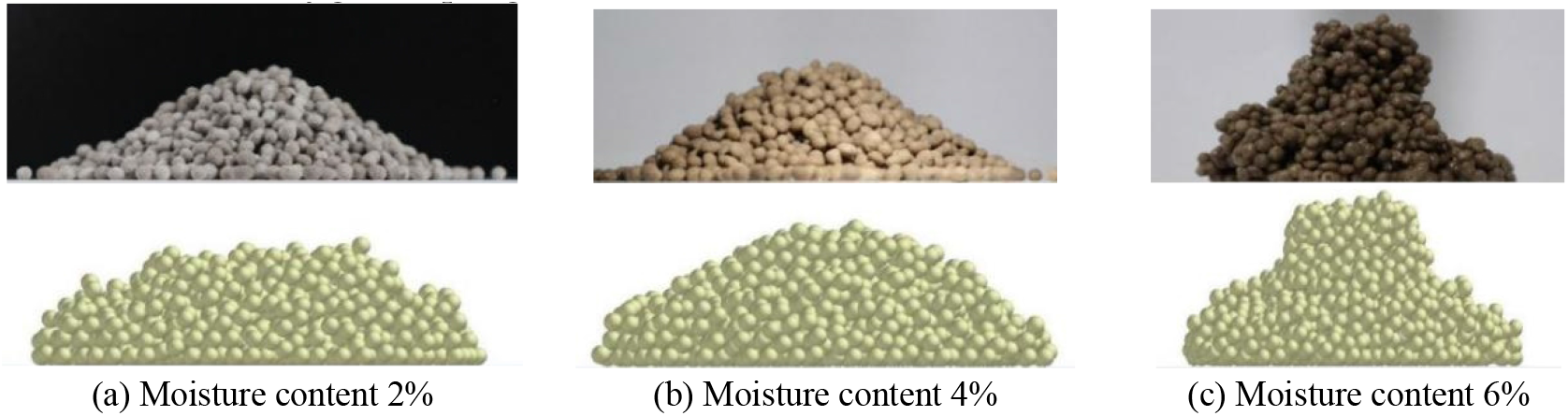
Validation of the angle of repose of a model fertilizer with moisture-absorbing granules at different moisture contents.

## 4. Moisture-absorbing viscous granular fertilizer discharge simulation and verification

To validate the precision of the model for moisture-absorbing granular fertilizers, simulations of a self-cleaning fertilizer device with a cam top plate were conducted, followed by bench validation experiments. These experiments aimed to compare and analyze the discharge error of the fertilizer in a single loop discharge simulation, with the performance of a single loop fertilizer discharge used as the evaluation criterion.

### 4.1 Moisture-absorbing viscous granular fertilizer EDEM-RecurDyn fertilizer discharge joint simulation

The cam top plate self-cleaning fertilizer discharge device depicted in Figure 6 operates by rotating the concave cam to push the fertilizer top plate, facilitating forced fertilizer discharge during the actual process. This movement cannot be replicated solely within the EDEM system. Hence, a collaborative effort between EDEM and the multi-body dynamics software RecurDyn was undertaken to construct a simulation model for the cam top plate self-cleaning fertilizer device. This model aimed to accurately simulate the fertilizer discharge process of the cam top plate self-cleaning fertilizer device.

**Fig. 6.**
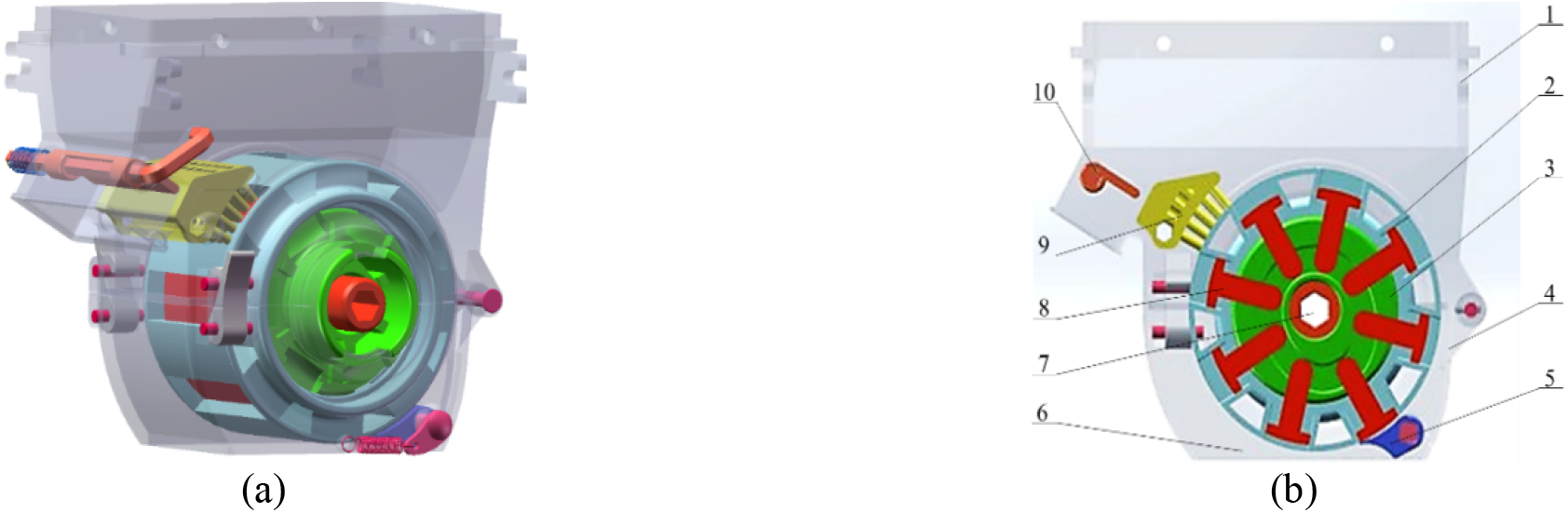
Structure diagram of cam top plate self-cleaning fertilizer device. (a:Three-dimensional model of fertilizer discharger, b:Sectional view of fertilizer discharger) 1. Upper housing 2. Feed unit wheel 3. Concave cam 4. Bottom shell 5. Fertilizer device 6. Ferilizer outlet 7. Intermdiate shaft 8. Fertilizer top plate 9. Fertilizer brush 10. Fertilizer unloading device.

The components of the cam top plate self-cleaning fertilizer device were imported into the RecurDyn multi-body dynamics software system in a 1:1 ratio using the step format. The material for each component of the fertilizer device was designated as PC board, rotational speed of the fertilizer device was configured to be 20 rad/min. The simulation model of the cam top plate self-cleaning fertilizer device was created by integrating the EDEM system with the RecurDyn system. A dynamic box was positioned directly beneath the fertilizer discharger to serve as a fertilizer collection container. The geometric model of the simulation is illustrated in Figure 7. The optimal parameter combinations for viscous granular fertilizer models with moisture contents of 2%, 4%, and 6% from Table 11 were incorporated into the fertilizer discharge simulation model. Following the fertilizer discharge simulation, the total simulation time was configured to be 10 s. In the EDEM post-processing module, the fertilizer discharge volume of a single loop under the stable fertilizer discharge state was determined. The simulation of fertilizer discharge for granular fertilizer with each moisture content was repeated three times.

**Fig. 7.**
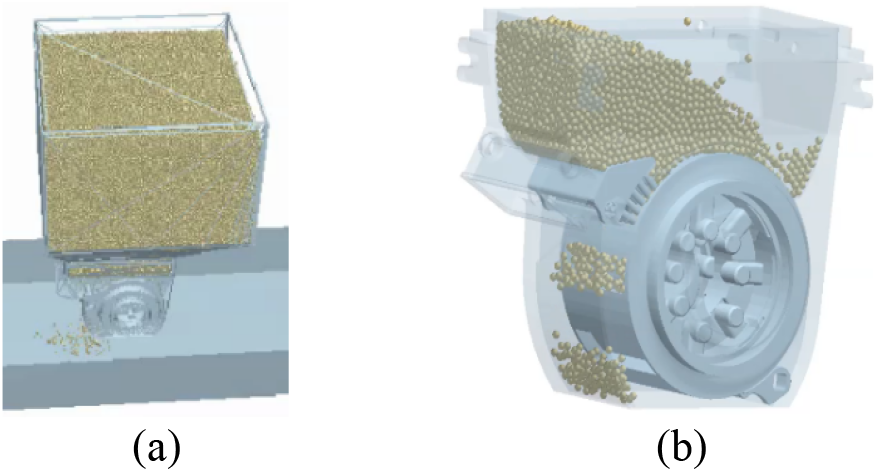
Simulation model of cam top plate self-cleaning fertilizer discharge. (a:Simulation model of cam top plate self-cleaning fertilizer discharge, b:Fertilizer diacharge process).

### 4.2 Bench validation experiment of moisture-absorbing viscous granular fertilizer

The equipment setup for the bench validation experiment comprises several components: a mounting bracket, fertilizer tank, fertilizer collection barrel, servo motor, machine governor, Hall speed sensor, and weighing system (Figure 8). Motor power is transmitted to the fertilizer discharge wheel via a six-square shaft and double-cross universal coupling. The Hall speed sensor was employed to oversee the rotational velocity of the fertilizer discharger. The fertilizer collection barrel and weighing sensor were positioned directly beneath the fertilizer outlet of the fertilizer discharger to promptly capture and measure the quantity of discharged fertilizer.

**Fig. 8.**
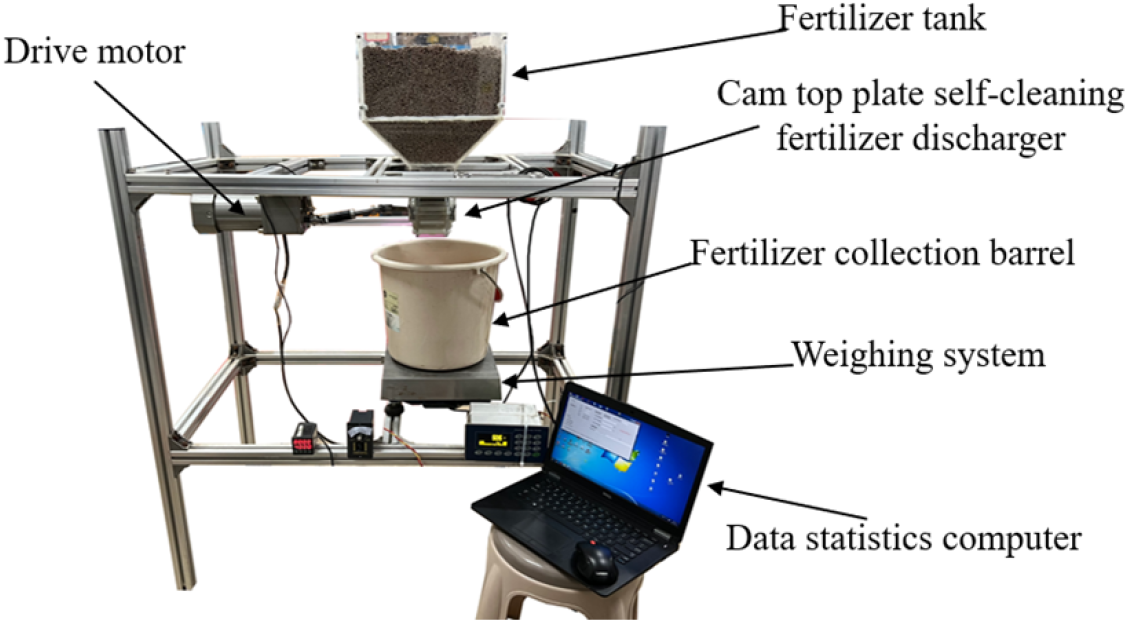
Actual fertilizer discharge test procedure

### 4.3 Results and analysis

Fig. 9 and Table 12 display the results of the fertilizer discharge simulation and bench validation experiments. The relative errors between the simulated single-loop fertilizer discharging volumes of moisture-absorbing viscous fertilizer with moisture contents of 2%, 4%, and 6%, and the actual single loop fertilizer discharging volume were 8.32%, 7.70%, 6.67%, respectively. The simulated single-loop discharging volume decreased with the increase in moisture content, mirroring the changing trend observed in the actual single-loop discharging volume. The reduction in discharging volume during the discharge process was attributed to the increase in moisture content, which led to enhanced viscosity and reduced mobility of the granular fertilizer.

**Table 12.** Rest angle results in simulations with different solution parameters for fertilizers with different moisture contents.

| Moisture | Stimulated single loop fertilizer discharging volume (g) | Actual single loop fertilizer discharging volume (g) | Relative error (%) |
| --- | --- | --- | --- |
| 2% | 48.58 | 52.99 | 8.32% |
| 4% | 46.74 | 50.64 | 7.70% |
| 6% | 39.07 | 41.86 | 6.67% |

**Fig. 9.**
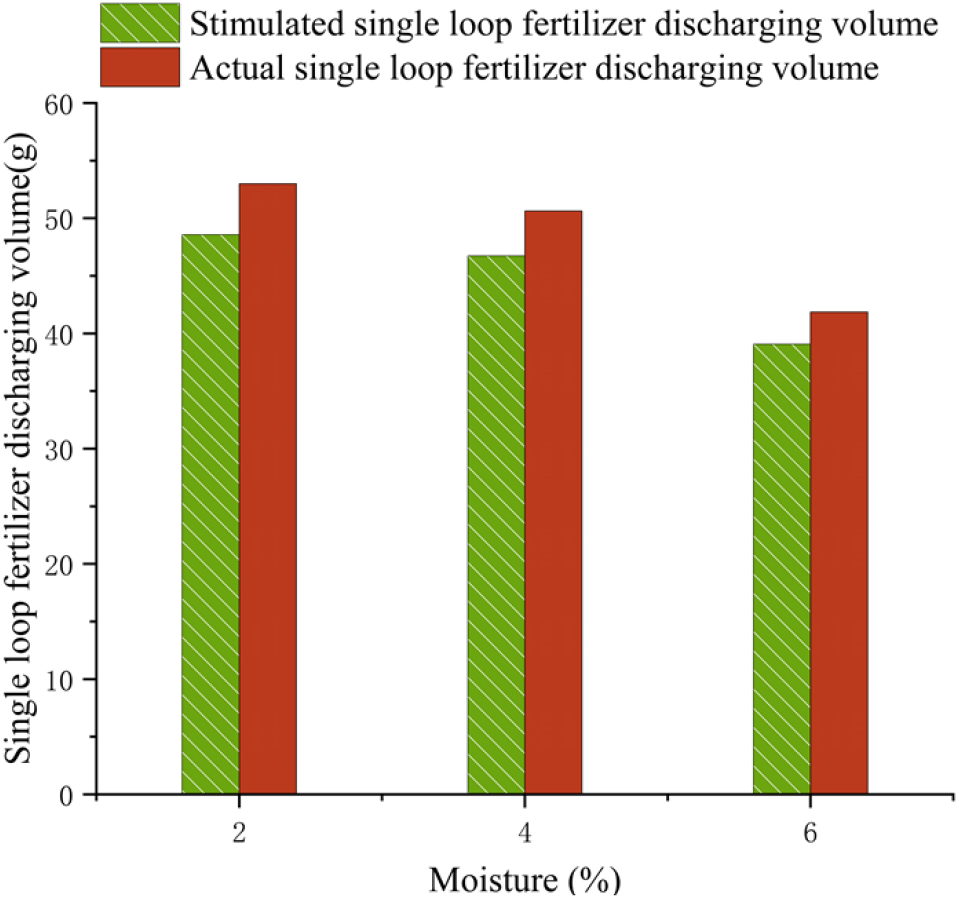
Percent bar graph of simulation and actual fertilizer discharge results

The aforementioned analysis indicates that the moisture content-discrete element significant parameter correlation model is capable of directly and precisely predicting the model parameters of moisture-absorbing viscous granular fertilizers across various moisture contents, this model can also be applied for constructing a discrete element model of moisture-absorbing viscous granular fertilizers considering their moisture absorption rate. The discrete element model of moisture-absorbing viscous granular fertilizers with varying moisture content accurately captures the viscous properties of these fertilizers and enables the simulation of interactions between moisture-absorbing viscous granular fertilizer and fertilizer discharger. This study offers research methodologies and theoretical models for analyzing the motion characteristics of viscous granular materials, as well as for the design and development of precision fertilizer discharging technology devices.

## 5. Conclusion

1. The discrete element parameter calibration was conducted for moisture-absorbing granular fertilizer with moisture contents ranging from 2% to 6%. This aimed to identify the significant parameters of the discrete element model specific to viscous granular fertilizer with 4% moisture content using Box Behnken test, Plackett Burman test, and hill-climbing test, with the optimal parameter combinations as fertilizer-fertilizer surface energy of 0.28 J/m^2^, fertilizer-PC complex coefficient of 0.21, and fertilizer shear modulus of 1.88×10^7^. The correlation model between significant parameters of the moisture-absorbing viscous fertilizer angle of rest and discrete elements was thoroughly rewritten, demonstrating a highly significant correlation with a P-value < 0.0001. The optimal parameter combination for discrete element model parameters of the viscous granular fertilizers with 2% and 6% moisture content was determined using correlation model between moisture-absorbing viscous granular fertilizer moisture content- and significant parameter correlation model.
2. The precision and dependability of the fertilizer’s moisture content-discrete element significant parameter correlation model were confirmed via cylinder lifting experiments, a comprehensive simulation and bench discharge experiment were conducted to simulate the fertilizer discharge process using the edem-recurdyn joint framework, the findings revealed that the relative error in the angle of rest prediction for the discrete element model of moisture-absorbing granular fertilizer, ranging from 2% to 6% moisture content, was ⩽1.42%, and stimulated single loop fertilizer’s relative error discharging moisture-absorbing granular fertilizer volume was ⩽ 8.32%. These results indicate that the moisture content-discrete element significant parameter correlation model is capable of accurately predicting the discrete element parameters of moisture-absorbing granular fertilizers across various moisture contents. Furthermore, the developed moisture-absorbing granular fertilizer model adeptly simulates surface interactions such as cohesion and mobility of viscous granular fertilizers. This capability furnishes valuable research methodologies and theoretical frameworks for scrutinizing the motion attributes of viscous granular materials, as well as for devising and refining precision fertilizer apparatus.

## Author Contributions

Chen Xiong Fei designed and conducted the simulation and experiment, and wrote the first draft of the article. Sun Ze Yu and Shi Yi Ze performed data analysis and image processing. Yu Jia Jia conducted literature review and commissioning of test equipment. Liu Jun An conceived and analyzed the data and drafted the article. Liu Mu Hua funded this research and provided all the experimental devices.

## Declaration of Competing Interest

The authors declare that they have no known competing financial interests or personal relationships that could have appeared to influence the work reported in this paper.

## Data availability

No data was used for the research described in the article.

## Funding

This work was funded by The National Natural Science Foundation of China (Grant No. 52165030) and Agricultural Machinery Equipment Application Industry Technology System of Jiangxi Province (Grant No. JXARS-21).

## Notes

### Competing Interest Statement

The authors have declared no competing interest.

